# Overnight memory consolidation facilitates rather than interferes with new learning of similar materials - a study probing NMDA-receptors

**DOI:** 10.1101/206771

**Authors:** Asfestani M Alizadeh, E Braganza, J Schwidetzky, J Santiago, S Soekadar, J Born, GB Feld

## Abstract

Whereas sleep-dependent consolidation and its neurochemical underpinnings have been strongly researched, less is known about how consolidation during sleep affects subsequent learning. Since sleep enhances memory, it can be expected to pro-actively interfere with learning after sleep, in particular of similar materials. This pro-active interference should be enhanced by substances that benefit consolidation during sleep, such as D-cycloserine. We tested this hypothesis in two groups (Sleep, Wake) of young healthy participants receiving on one occasion D-cycloserine (175 mg) and on another occasion placebo, according to a double-blind balanced cross-over design. Treatment was administered after participants had learned a set of word-pairs (A-B list) and before nocturnal retention periods of sleep vs. wakefulness. After D-cycloserine blood plasma levels had dropped to negligible amounts, i.e., the next day in the evening, participants learned, in three sequential runs, new sets of word-pairs. One list – to enhance interference – consisted of the same cue words as the original set paired with a new target word (A-C list) and the other of completely new cue words (D-E set). Unexpectedly, during post-retention learning the A-C interference list was generally better learned than the completely new D-E list, which suggests that consolidation of previously encoded similar material enhances memory integration rather than pro-active interference. Consistent with this view, new learning of word-pairs was better after sleep than wakefulness. Similarly, D-cycloserine generally enhanced learning of new word-pairs, compared to placebo. This effect being independent of sleep or wakefulness, leads us to speculate that D-cycloserine, in addition to enhancing sleep-dependent consolidation, might mediate a time-dependent process of active forgetting.

## Introduction

The relationship between sleep and memory maintenance has received detailed attention in the last 20 years (Diekelmann & Born, 2010; Rasch & Born, 2013) and there is widespread interest in enhancing this beneficial effect of sleep on memory (Feld & Diekelmann, 2015), e.g., by enhancing neuronal oscillations (Marshall, Molle, Hallschmid, & Born, 2004; Ngo, Martinetz, Born, & Molle, 2013) or externally cueing replay (Rasch, Buchel, Gais, & Born, 2007; Rudoy, Voss, Westerberg, & Paller, 2009) i.e., processes that support sleep-dependent memory. We recently demonstrated that the N-methyl-D-aspartate (NMDA) receptor co-agonist D-cycloserine powerfully enhances sleep-dependent declarative memory consolidation when administered before sleep (Feld, Lange, Gais, & Born, 2013). It is however completely unclear, how this enhancement affects the subtle balance of encoding and memory maintenance in the brain (Richards & Frankland, 2017). This is especially interesting as sleep has been suggested to also benefit new learning (Mander, Santhanam, Saletin, & Walker, 2011).

One of the first reports investigating the effect of sleep on memory was by Jenkins and Dallenbach (1924), who famously hypothesised that sleep enhances memory not via an active process but by shielding it from interference, a line of argument that remains popular (Mednick, Cai, Shuman, Anagnostaras, & Wixted, 2011). However, since this proposal it has been convincingly shown across species, modalities and paradigms that during sleep memory is actively strengthened by the repeated replay of traces that were encoded during prior phases of wakefulness (Rasch et al., 2007; Rudoy et al., 2009; Sadowski, Jones, & Mellor, 2016; van de Ven, Trouche, McNamara, Allen, & Dupret, 2016; Wilson & McNaughton, 1994). Intriguingly, it has also been shown that this sleep-dependent consolidations makes memory traces more robust towards retro-active interference (Ellenbogen, Hulbert, Stickgold, Dinges, & Thompson-Schill, 2006), i.e.,, to the interfering influence of learning new information that deteriorates the original trace even if it had previously been successfully encoded (Osgood, 1948). When participants in this experiment learned a set of word-pairs (A-B) before sleep and had to learn a retro-actively interfering set of word-pairs (A-C) after sleep (but before retrieval), the effect of sleep on memory retention was enhanced. Moreover, in a study where participants encoded while exposed to the smell of roses, re-exposing them to this odour cue during sleep made the associated memory robust to retro-active interference and the same treatment during wakefulness had the opposite effect (Diekelmann, Wilhelm, Wagner, & Born, 2011). These findings pose the intriguing question whether the reduction in memories’ susceptibility to retro-active interference during sleep is due to a strengthening of the original trace that would be accompanied by enhanced pro-active interference, i.e., whether new memory traces are harder to establish if they overlap with these stronger old memory traces (Osgood, 1948)

The oscillatory properties of sleep that support the consolidation process (Staresina et al., 2015) are ideally suited to drive the strengthening of memory traces via long term potentiation (LTP) (King, Henze, Leinekugel, & Buzsaki, 1999), which occurs mainly at glutamatergic synapses and is mediated by NMDA receptors (Malenka & Bear, 2004; Malenka & Nicoll, 1999). Accordingly, we administered D-cycloserine, a drug that supports NMDA receptor activation by binding to its glycine binding site (Sheinin, Shavit, & Benveniste, 2001), to participants after they learned word-pairs, so that peak plasma concentration occurred during the first half of the sleep phase (Feld et al., 2013). Enhancing NMDA receptor activation benefitted the sleep-dependent consolidation specifically of the word-pairs if given during sleep and thus represents the ideal model to test whether memory traces enhanced by sleep introduce detrimental pro-active interference on new learning.

To test this we asked participants to learn a list of word-pairs (A-B) and then enhanced sleep-dependent consolidation of these memories by administering D-cycloserine (Feld et al., 2013). We expected that, when participants learned a new list of word-pairs (AC) the next evening (i.e., after twice the drug half-life), performance would be reduced under treatment compared to placebo due to enhanced pro-active interference of the more strongly consolidated memory. To specify whether this effect depends on the item specificity of proactive interference, participants also learned new word-pairs that did not overlap with the original list (D-E) and we expected that performance on this list would not be affected by treatment. We also tested a group of participants that did not sleep during the retention interval to test our hypothesis that these effects are mediated by processes active only during sleep.

## Methods

### Participants

Fifty-one participants completed the study in the sleep (n = 30) and the Wake (n = 21) conditions. Participants were healthy, non-smoking, native German speaking men, age range between 18-30 years old, with a body mass index between 19-26 kg/m^2^. Before starting the study a routine medical examination was performed for all the participants to exclude any psychiatric, neurological or endocrine diseases, participants who took regular medication were also excluded. The medical examination relied on a structured interview asking for current or past diagnosed conditions as well as a blood pressure and blood screening test. Participants were not allowed to take any acute medication at the time of the experiments and they reported a normal sleep-wake cycle and no shift work for at least 6 weeks before the experiments. They were instructed to get up at 07:00 h on experimental days and, during these days, not to take any naps, no caffeine-containing drinks after 13:00 h and also not to consume alcohol one day before the experimental nights. Before the sleep experiment, participants took part in an adaption night under conditions of the experiment (i.e., including the placement of electrodes for polysomnographic recordings). The experiments were approved by the ethics committee of the University of Tübingen. We obtained written informed consent from all participants before their participation.

### Design and procedures

The experiments followed a double-blind, placebo-controlled, within-subject crossover design, with Sleep vs. Wake between-subjects groups. Within each group, participants took part in two identical experimental sessions with the exception of administration of D-cycloserine or placebo (D-cycloserine: Cycloserine Capsules®, 175 mg, the Chao Center for Industrial Pharmacy & Contract Manufacturing, USA, plasma halftime: 10 h, plasma maximum: 1-2 h; Figure 1 summarizes study design). The two experimental sessions were scheduled at least 14 days apart.

**Figure 1:**
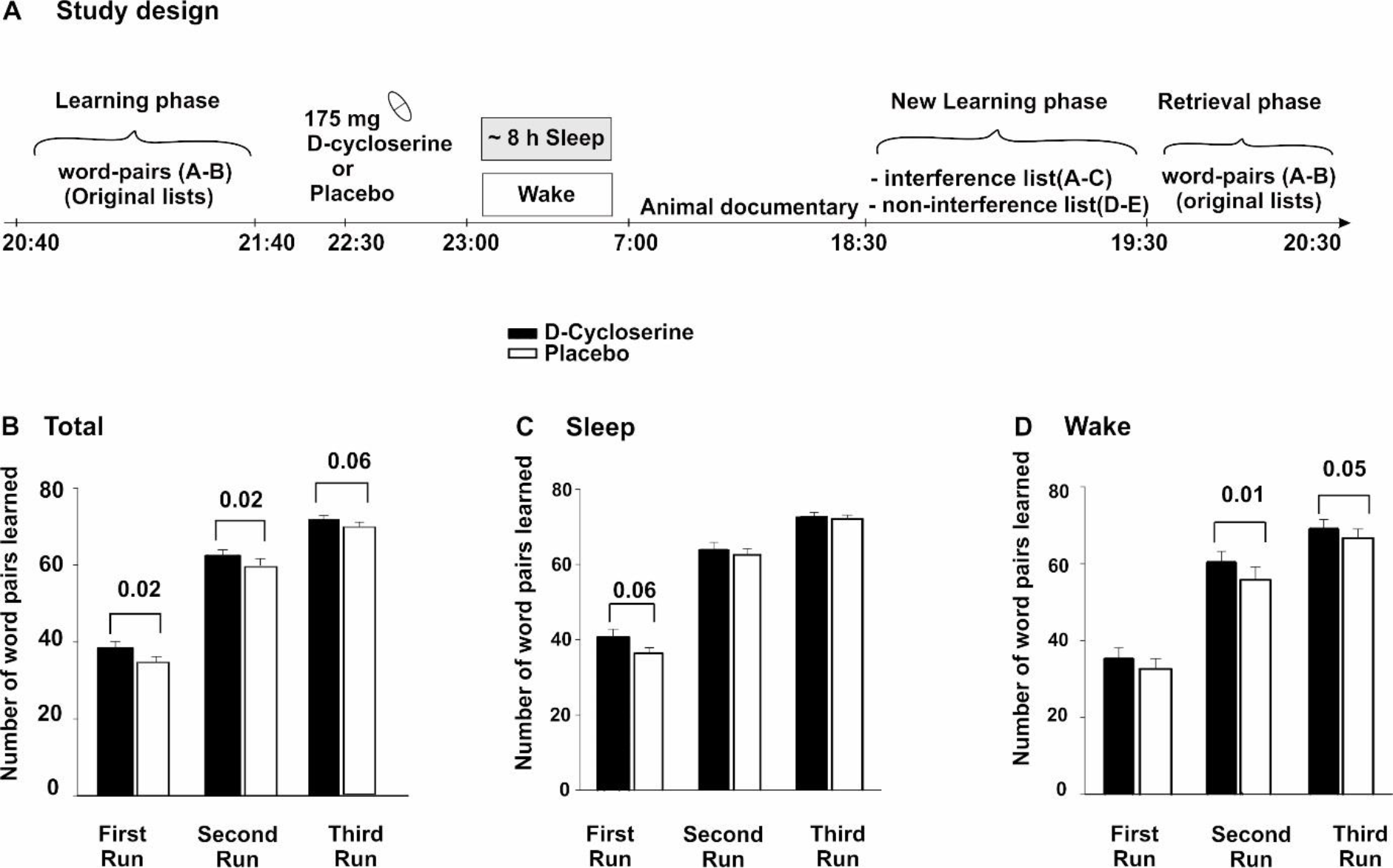
(A) Participants first learned 80 word-pairs (A-B) up to a criterion of 60%, by a repeated cued recall procedure (see Methods section for details). Afterwards at approximately 22:30 h, they took 175 mg D-cycloserine or placebo. At 23:00 h the participants in the Sleep group went to bed and polysomnographic recording was performed, whereas the Wake participants watched documentaries about planets. All participants received breakfast at 7:45 h and watched animal documentaries until 18:00 h. Afterwards, at 18:30 h, the participants learned 80 new word-pairs in three consecutive runs and finally retrieved the original 80 word-pairs. (B) Mean and standard error of the mean (SEM) of the amount of correctly recalled word-pairs in in total, (C) in the Sleep experiment and (D) in the Wake experiment during the three runs of the New Learning phase are shown.

**Figure 2.**
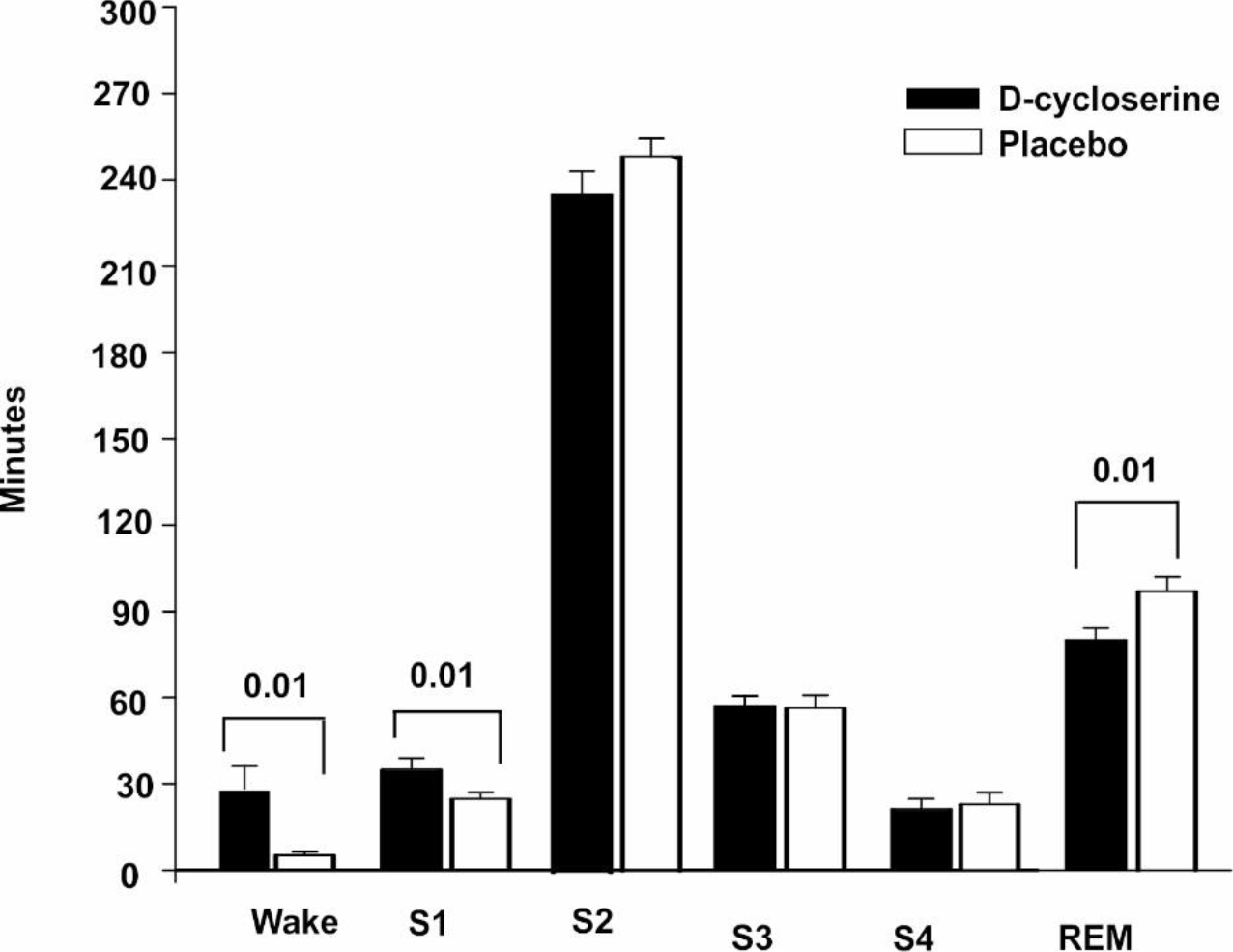
Mean (SEM) time spent in the different sleep stages Wake, S1 (sleep stage 1), S2 (sleep stage 2), SWS (slow wave sleep) and REM (rapid eye movement sleep) in minutes.

Participants arrived at 20:00 h for each experimental night and first filled out questionnaires. In the Sleep group, polysomnography was prepared by applying electrodes. Next they learned the first set of declarative word-pairs (original word-pairs) between 20:40 h and 21:40 h. Participants were informed that the word-pairs would be recalled immediately and also the next evening, as well as, that they would learn new word-pairs during the next evening. After learning they filled in questionnaires measuring mood and sleepiness and performed a reaction time task to measure vigilance (psychomotor vigilance task). At 22:30h, Participants received the medication (D-cycloserine or placebo). At 23:00 h, the electrodes were connected to the amplifiers and lights were turned off, in the Sleep group. The Wake group watched astronomy documentaries (two counter balanced lists of films, one for each session) in the lab during this time. After approximately 8 hours (between 6:45 h and 7:15 h), the Sleep group was woken up (if possible from sleep stage 1 or 2). All participants first answered questionnaires measuring their mood and sleepiness and, afterwards, received a standardised breakfast consisting of bread, cheese and/or salami, honey and jam and, herbal tea (no caffeine) and the Sleep group was allowed to shower to clean the electrode gel off their head. During the day, participants followed a tight protocol watching animal documentaries (two counter balanced lists of films, one for each session) for approximately one and half hours at a time (two episodes), followed by a break to take a walk around the campus and, received two snacks in the afternoons. One snack consisted of fruit tea, biscuits and an apple, the other one was two slices of bread, cheese and salami, fruit tee, tomato and, butter. This was done for 10 hours only interrupted by lunch at the local canteen together with the investigator. This protocol was chosen to standardize the experience of the participants after drug application, which should include as little as possible the opportunity of newly encoding written words. At 18:30 h, participants learned new word-pairs and immediately recalled them. Next they recalled original word-pairs they learned the day before. Finally, we again measured mood, sleepiness and vigilance, as well as, word generation.

### Word-pair tasks

Consolidation was measured using 80 slightly associated word-pairs (A-B) in two lists (original lists). The word-pairs were presented on a computer screen for 4 seconds each with a 1 second inter-stimulus interval (ISI). After presenting both lists, the participant’s memory was tested in a cued recall procedure by presenting only the first word and asking the participant to produce the associated word. This was done for each list individually. If the participant did not reach the criterion of 60% correct responses on one (or both) of the lists, only this list (or both lists) was presented again completely (3 seconds per pair) and cued recall was repeated. This was done until he reached the criterion. The amount of word-pairs recalled during the last cued recall was used as measure of learning performance. The cued recall procedure as described above was performed again at the very end of the Retrieval phase (after the New Learning phase – see below). This was done to keep the participants motivated to consolidate the memory traces across both sessions, which has previously been shown to be an essential factor driving sleep-dependent memory consolidation (Wilhelm et al. 2011); data from this retrieval were not analysed as they are confounded by the prior new learning).

During the next evening, participants were presented a new set of 80 word-pairs in two separate lists. One list with forty word-pairs (A-C list) interfered with the original list, i.e., they contained a new second word (C) paired with a cue word (A) of the original list (interference list), the other also consisting of forty word-pairs (D-E list) was completely new (no-interference list). These two word-pair lists were also learned back to back in a balanced order and each pair was shown for 4 seconds (1 second ISI). The cued recall procedure described above was performed three times (run 1 - run 3) and after runs 1 and 2 the word-pairs were shown again for 3 seconds each.

### Polysomnography, sleep analysis, and EEG power analysis

The EEG was recorded continuously from electrodes (Ag-AgCl) placed according to the 10-20 System, referenced to two linked electrodes attached to the mastoids. EEG signals were filtered between 0.03 and 35 Hz and sampled at a rate of 250 Hz using a Brain Amp DC (BrainProducts GmbH, Munich, Germany). Additionally, horizontal and vertical eye movements (HEOG, VEOG) and the EMG (via electrodes attached to the chin) were recorded for standard polysomnography. Sleep architecture was determined according to standard polysomnographic criteria using EEG recordings from C3 and C4 (Rechtschaffen and Kales, 1968). Scoring was performed by an experienced technician who was blind to the assigned treatment (an additional expert was consulted for ambiguous epochs). For each night, total sleep time (TST), i.e., the time between the first detection of transition from sleep stage 1 to 2 and lights on, and time spent in the different sleep stages, i.e., wake; sleep stages 1, 2, 3, 4; SWS (defined by the sum of sleep stage 3 and 4) and rapid eye movement (REM) sleep was calculated in minutes.

### Control measures - vigilance, sleepiness, and mood ratings and test of encoding

Participant’s sleepiness and mood was assessed using self-report measures. The Stanford Sleepiness Scale (SSS) (Hoddes, Zarcone, Smythe, Phillips, & Dement, 1973) measures subjective sleepiness with one item and eight answer options ranging from one = “Feeling active, vital, alert, or wide awake” to eight = “Asleep” (provided as an anchor). We assessed the participant’s mood using the multidimensional mood questionnaire at three time points per session (Hinz, Daig, Petrowski, & Brahler, 2012). This questionnaire produces the three scales positive mood (high is positive), tiredness (low is tired) and calmness (high is calm). Objective vigilance was additionally tested using the psychomotor vigilance task (PVT; (Dinges et al., 1997)). This 5-min version of the PVT required pressing a button as fast as possible whenever a bright millisecond clock presented on a dark computer screen started counting upward. After the button press, this clock displayed the reaction time. General capabilities of long-term memory retrieval were tested using a word generation task Participants had to produce as many words as possible starting with a certain letter (P or M) or belonging to a defined category (hobby or profession) during a time of 2 minutes each (Regensburger Wortflüssigkeitstest [WFT]; Aschenbrenner et al., 2000). At the end of each session all participants were asked if they believed to have received the active agent or placebo.

### Data reduction and statistical analysis

In the Sleep group two participants were excluded because of their extremely low learning performance (below 7 word-pairs in more than one list) and two participants were excluded because of disrupted sleep. Statistical analyses generally relied on analyses of variance (GLM; SPSS version 21.0.0 for Windows) including the repeated-measures factors Substance (D-cycloserine vs placebo), Interference (interference vs no-interference) and, where appropriate, the factor Runs (1,2,3) pertaining to the three recalls during the New Learning phase, as well as, the between-subjects factor Sleep/Wake. Greenhouse-Geisser correction of degrees of freedom was applied where necessary. Significant interactions were followed up by lower level ANOVAs and post-hoc t-tests.

### Results

#### Word*-pairs*

The interference list was learned significantly better than the no-interference list, with this effect being predominant on the first run (Interference × Run: F_(2,90)_ = 9.035, p = 0.001, First run: F_(1,45)_ = 9.492, p = 0.004; Table 1). Trivially, performance improved across the three runs (F_(2,90)_ = 832.695, p ≤ 0.001). We also found a trend towards the Sleep group learning more new word-pairs than the Wake group (F_(1,45)_ = 3.447, p = 0.070, for main effect of the Sleep/Wake factor). D-cycloserine distinctly enhanced new learning of word-pairs 20 hours after administration (F_(1,45)_ = 6.512, p = 0.014, for main effect of Substance). On average participants learned 2.7 more new word-pairs in the D-cycloserine condition than in the Placebo condition (D-cycloserine: 57.30 ±1.40, Placebo: 54.63 ±1.43).

We also identified a three-way interaction of Substance*Run*Sleep/Wake (F_(2,90)_ = 3.514, p = 0.034). This effect was mainly driven by word-pairs being learned more in the Sleep group after D-cycloserine during the first run (t_(25)_ = 1.964, p = 0.061), whereas word-pairs being learned more in the Wake group after D-cycloserine during the second (t_(20)_ = 2.880, p = 0.009) and third (t_(20)_ = 2.102, p = 0.048) runs. All the other effects and interactions were non-significant (All p ≥ 0.114; Table 1).

**Table 1:** Mean (SEM) correctly recalled word-pairs in the New Learning phase for the interference and no-interference conditions.

|  | Sleep |  |  |  | Wake |  |  |  |
| --- | --- | --- | --- | --- | --- | --- | --- | --- |
|  | D-cycloserine |  | Placebo |  | D-cycloserine |  | Placebo |  |
| **Interference** |  |  |  |  |  |  |  |  |
| **First run** | 21.42 | (1.12) | 19.00 | (0.93) | 19.38 | (1.32) | 16.67 | (1.49) |
| **Second run** | 31.85 | (0.82) | 31.27 | (1.00) | 30.76 | (1.25) | 27.57 | (1.57) |
| **Third run** | 36.08 | (0.53) | 36.31 | (0.59) | 35.10 | (1.13) | 33.00 | (1.22) |
| **No-Interference** |  |  |  |  |  |  |  |  |
| **First run** | 19.27 | (1.28) | 17.38 | (0.98) | 15.95 | (1.61) | 16.00 | (1.62) |
| **Second run** | 32.23 | (1.20) | 31.31 | (0.99) | 29.67 | (1.62) | 28.19 | (1.97) |
| **Third run** | 36.73 | (0.62) | 35.81 | (0.65) | 34.00 | (1.34) | 33.57 | (1.26) |

#### Sleep stages

Under D-cycloserine participants spent significantly more time (in minutes) in wakefulness and sleep stage 1 (Wake: t_(25)_ = −2.737 p = 0.011; stage 1: t_(25)_ = −2.661 p = 0.013) and less time in REM sleep (t_(25)_ = 2.768, p = 0.010). We also found a trend towards reduced time in sleep stage 2 in the D-cycloserine condition in comparison to placebo (t_(25)_ = 1.795, p = 0.085) but there was no significant difference between the treatments in time spent in the other sleep stages (all t ≥ −0.356, p ≥ 0.451) or total sleep time (t_(25)_ = −0.509, p = 0.615).

#### Control measures

In both the Sleep and Wake group we found no differences between the treatments in the psychomotor vigilance task (PVT, all p ≥ 0.196). In the Sleep group, subjective ‘tiredness’ in the morning after nocturnal sleep, was enhanced in the D-cycloserine group (t_(25)_ = −2.534, p = 0.018; Table 2). In the Wake group, subjects in the morning after D-cycloserine showed trend-wise higher ‘good mood’ and less ‘tiredness’ (‘tiredness’: t_(20)_ = 1.910, p = 0.071, ‘good mood’: t_(20)_ = 1.805, p = 0.086; Table 2) than after placebo. Sleepiness (on the SSS did not differ between groups and substance conditions at all times (all p ≥ 0.167). Also, we did not have any significant differences between groups and substance conditions in the general retrieval performance as measured by the word fluency task (all p ≥ 0.503). Participants in both groups were not able to discriminate between D-cycloserine and placebo (McNemars’ exact test: p ≥ 0.774).

**Table 2:**
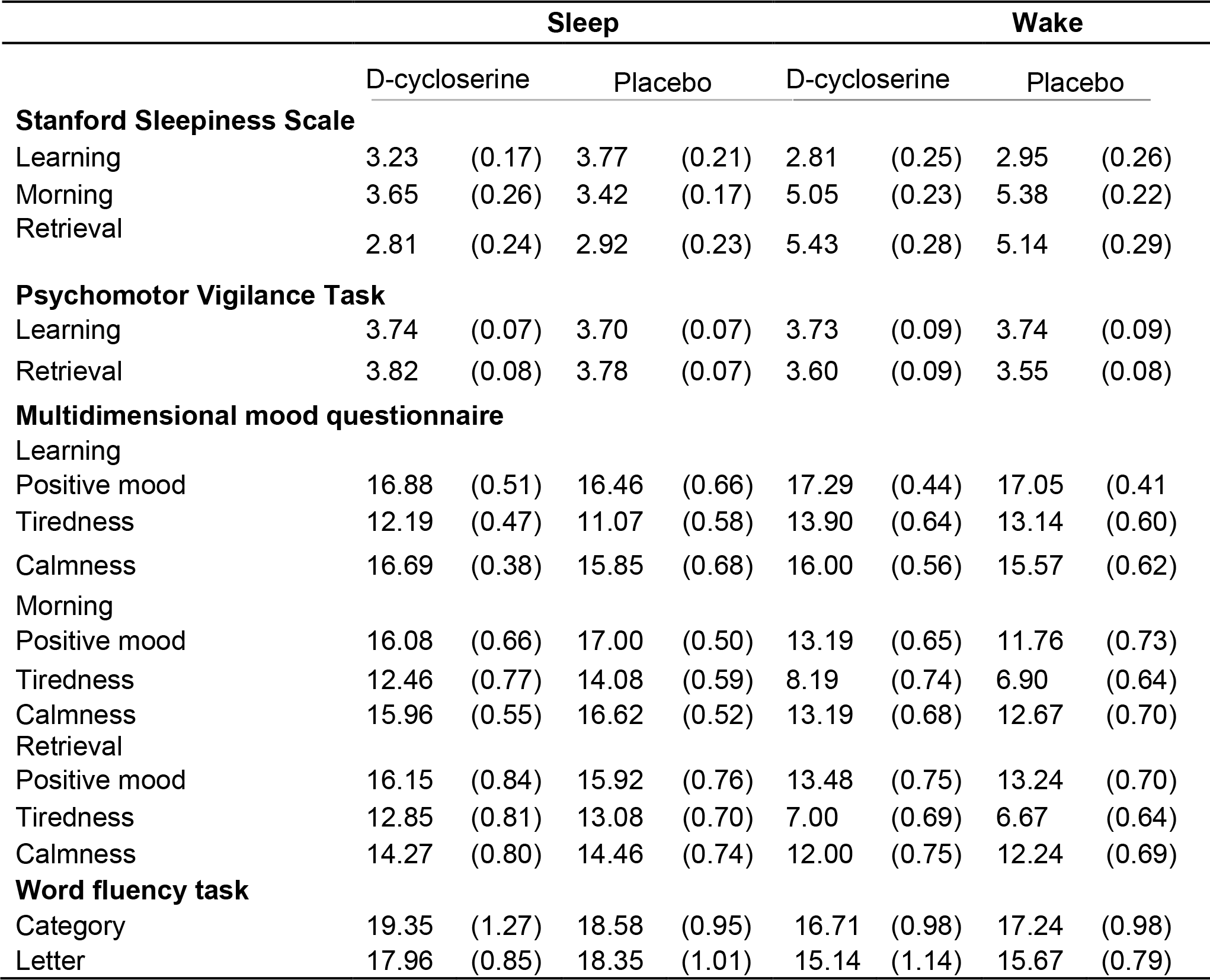
Means (SEM) of performance are given for the **c**ontrol measures. Subjective sleepiness (Stanford Sleepiness Scale), objective vigilance (Psychomotor Vigilance Task), mood (Multidimensional mood questionnaire) and general retrieval performance (Word Fluency Test). Learning (after the original Learning phase), Morning (at approximately 7:15 h), Retrieval (after the Retrieval phase).

|  | Sleep |  |  |  | Wake |  |  |  |
| --- | --- | --- | --- | --- | --- | --- | --- | --- |
|  | D-cycloserine |  | Placebo |  | D-cycloserine |  | Placebo |  |
| Stanford Sleepiness Scale |  |  |  |  |  |  |  |  |
| Learning | 3.23 | (0.17) | 3.77 | (0.21) | 2.81 | (0.25) | 2.95 | (0.26) |
| Morning | 3.65 | (0.26) | 3.42 | (0.17) | 5.05 | (0.23) | 5.38 | (0.22) |
| Retrieval | 2.81 | (0.24) | 2.92 | (0.23) | 5.43 | (0.28) | 5.14 | (0.29) |
| Psychomotor Vigilance Task |  |  |  |  |  |  |  |  |
| Learning | 3.74 | (0.07) | 3.70 | (0.07) | 3.73 | (0.09) | 3.74 | (0.09) |
| Retrieval | 3.82 | (0.08) | 3.78 | (0.07) | 3.60 | (0.09) | 3.55 | (0.08) |
| Multidimensional mood questionnaire |  |  |  |  |  |  |  |  |
| Learning |  |  |  |  |  |  |  |  |
| Positive mood | 16.88 | (0.51) | 16.46 | (0.66) | 17.29 | (0.44) | 17.05 | (0.41) |
| Tiredness | 12.19 | (0.47) | 11.07 | (0.58) | 13.90 | (0.64) | 13.14 | (0.60) |
| Calmness | 16.69 | (0.38) | 15.85 | (0.68) | 16.00 | (0.56) | 15.57 | (0.62) |
| Morning |  |  |  |  |  |  |  |  |
| Positive mood | 16.08 | (0.66) | 17.00 | (0.50) | 13.19 | (0.65) | 11.76 | (0.73) |
| Tiredness | 12.46 | (0.77) | 14.08 | (0.59) | 8.19 | (0.74) | 6.90 | (0.64) |
| Calmness | 15.96 | (0.55) | 16.62 | (0.52) | 13.19 | (0.68) | 12.67 | (0.70) |
| Retrieval |  |  |  |  |  |  |  |  |
| Positive mood | 16.15 | (0.84) | 15.92 | (0.76) | 13.48 | (0.75) | 13.24 | (0.70) |
| Tiredness | 12.85 | (0.81) | 13.08 | (0.70) | 7.00 | (0.69) | 6.67 | (0.64) |
| Calmness | 14.27 | (0.80) | 14.46 | (0.74) | 12.00 | (0.75) | 12.24 | (0.69) |
| Word fluency task |  |  |  |  |  |  |  |  |
| Category | 19.35 | (1.27) | 18.58 | (0.95) | 16.71 | (0.98) | 17.24 | (0.98) |
| Letter | 17.96 | (0.85) | 18.35 | (1.01) | 15.14 | (1.14) | 15.67 | (0.79) |

## Discussion

We expected that learning of new word-pairs will be decreased after sleep under D-cycloserine in comparison to placebo, due to enhanced pro-active interference by the consolidated memory. This effect was predicted to be facilitated, if the new word-pairs shared the consolidated word-pair’s cue word. In contrast, we found that sharing a cue word with the original list enhanced new learning rather than impaired it. Also, our results provide evidence that new learning was generally facilitated by sleep and by D-cycloserine. Notably the enhancing effect of D-cycloserine was independent of whether it was given before sleep or wakefulness, which was also unexpected. The effects of D-cycloserine on sleep architecture replicated earlier findings (Feld et al., 2013). Generally, these findings suggest that pro-active interference that is predominant immediately after learning does not carry over to longer retention intervals but rather is reversed by consolidation to aid new learning. Accordingly, enhancing sleep-dependent consolidation of memory traces appears to proactively support new learning. Moreover, the effect of D-cycloserine being independent of sleep and associated consolidation, suggests additional time-dependent mechanisms supporting new learning perhaps by inducing active forgetting.

Our finding of better encoding in the interference condition than in the no-interference condition indicates that rather than producing pro-active interference and thereby impairing new learning our interference condition enhanced new learning. This cannot be explained by the initial memory merely decaying across time and thereby reducing its pro-active influence, as this would not enhance performance on the interference above and beyond the no-interference condition. It has been proposed that new information can be learned more easily, if it can be integrated with existing knowledge (van Kesteren, Rijpkema, Ruiter, Morris, & Fernandez, 2014). Theoretically consolidation during sleep may derive such knowledge by abstracting from episodes (Lewis & Durrant, 2011). However, empiric evidence suggests that knowledge may be built in a time-rather than a sleep-dependent manner (Hennies, Lewis, Durrant, Cousins, & Ralph, 2014), which is consistent with our data revealing that improved learning of interfering materials is independent of prior sleep or wakefulness. Essentially, this question needs to be addressed by additional experiments that go beyond the scope of the current study and establish when pro-active interference is overridden by knowledge abstraction.

The trend-wise enhanced encoding in the Sleep versus the Wake group, corresponds to earlier findings of enhanced encoding after sleep (Mander et al., 2011). Interestingly, we found that there was a significant three-way interaction (between Substance, Sleep/Wake, and Runs), which appeared to mainly reflect that in the Sleep group D-cycloserine enhanced new learning on run 1, whereas in the Wake group the NMDA-receptor co-agonist enhanced learning on runs 2 and 3. This can be interpreted as the Sleep group already reaching ceiling levels after run 1, because of the additional boost in learning through sleep. The Wake group on the other hand had more opportunity to increase learning later on in the task. In essence, we suggest that this interaction effect is mediated by sleep-dependent increases in new learning that are independent of the increases in new learning induced by D-cyloserine.

Thus, unexpectedly, D-cycloserine not only enhanced new learning when administered before sleep but also when administered before a wake retention period. This is difficult to integrate. We suspect that this effect might reflect an involvement of NMDA-receptor activation in sleep-independent processes that renormalize synaptic weights and generally free capacity for novel encoding (see (Tononi & Cirelli, 2014) and (Frank, 2012) for opposing remarks) that have recently been shown to also occur during wakefulness (Hengen, Torrado Pacheco, McGregor, Van Hooser, & Turrigiano, 2016). Active decay is a form of renormalisation that has been suggested to occur at glutamatergic synapses (Hardt, Nader, & Nadel, 2013). Studies of object-location and associative memories in rats have shown that such decay can be prevented by blocking the removal of α-amino-3-hydroxy-5-methyl-4-isoxazolepropionic acid (AMPA) receptors from the synapse (Migues et al., 2016). Similarly, blocking NMDA-receptors for prolonged periods impaired forgetting of spatial memory in rats (Villarreal, Do, Haddad, & Derrick, 2002). As activating the NMDA-receptor in specific ways induces AMPA-receptor endocytosis (Beattie et al., 2000), it is tempting to speculate that the present finding of a generally enhanced encoding (after wake as well as sleep retention periods) involves D-cycloserine sensitising NMDA-receptors to ambient glutamate levels (Featherstone & Shippy, 2008), which drives forgetting and frees up capacity for new learning. It is important to note that D-cycloserine has also been shown to directly enhance performance when administered before learning (Ledgerwood, Richardson, & Cranney, 2003), and although unlikely we cannot exclude that, after two times the half-life of the drug, a residual direct influence on new learning remained.

In conclusion, we found that overnight retention periods after learning facilitated new learning in particular of interfering materials. Sleep as well as D-cycloserine generally enhanced new learning, and these effects might partly originate from their consolidating influence of the originally learned A-B word pairs, that might facilitate transfer learning of new word-lists (including A-C and D-E lists). The effect of D-cycloserine likewise observed after wake periods also suggests a contribution of NMDA-receptor mediated active decay (Hardt et al., 2013) that is established as a form of sleep-independent synaptic renormalisation (Frank, 2012; Hengen et al., 2016). Examining how this form of forgetting interacts with sleep-dependent forms of synaptic renormalisation (Tononi & Cirelli, 2014) and sleep-dependent memory consolidation (Diekelmann & Born, 2010), will be of essence to understand how consolidation and forgetting sustain long-term memory and new learning (Feld & Born, 2017).

## Acknowledgements

The authors gratefully acknowledge Marion Inostroza for supporting all phases of the project and also would like to thank Michael Radloff for assisting data collection.

